# Epigenetic Profiling Provides Insights into AIDS Resistance in African Green Monkey

**DOI:** 10.1101/535567

**Authors:** Zheng Yuan, Xiao-jun Zhou

**Affiliations:** Laboratory Animal Center of the Academy of Military Medical Sciences, Beijing, China

## Abstract

As a natural host of simian immunodeficiency virus (SIV), African green monkeys (AGM) do not develop AIDS although high levels of SIV replication were maintained. Low frequencies of CD4^+^ T cells and high frequencies of CD8^dim^ T cells were observed in healthy adult AGM, which may partially explain the absence of SIV-induced disease progression. Elucidating the mechanisms that allow this natural host co-exist with SIV without progressive disease may facilitate knowledge of AIDS pathogenesis. Here we show: (1) Compared with junior AGM, 3 miRNA were up-regulated in adult AGM in which hsa-miR-151a-3p was shown to target both *CD4* and *MAZR*; 15 miRNAs were down-regulated in adult AGM in which hsa-miR-140-5p, hsa-miR-126-3p and hsa-miR-194-5p were shown to target *CD8α;* (2) MeDIP sequencing analysis of adult AGM samples revealed that hypermethylation exists in the promoter region of *CD4*, *CXCR6, CCR5*, while hypomethylation exists in the promoter region of *RUNX3*, *ICAM2*; (3) Hypomethylation in the promoter region of *PTK2* contributes to up-regulated expression of hsa-miR-151a-3p in adult AGM, while hypermethylation in the promoter region of *WWP2* contributes to down-regulated expression of hsa-miR-140-5p. Our data for the first time demonstrates the link between miRNA and DNA methylation expression profile, which may together contribute to the phenotype of AIDS resistance in AGM.

**Author Summary:** African green monkeys (AGM) do not develop AIDS although high levels of SIV replication were maintained. Elucidating the mechanisms that allow this natural host co-exist with SIV without progressive disease may facilitate knowledge of AIDS pathogenesis. In this study, the miRNA expression patterns were found to be associated with the switch from CD4^+^ to CD4^−^CD8a^dim^ in adult AGM. The up-regulated hsa-miR-151a-3p was shown to target both AGM *CD4* and *MAZR*, while the down-regulated hsa-miR-140-5p, hsa-miR-126-3p and hsa-miR-194-5p were shown to target AGM *CD8α*. And none of these miRNAs possess target sites in cynomolgus macaque (CM) *CD4, CD8α* and *MAZR* reflecting differences in AIDS resistance between these two species. Our data also demonstrates the link between miRNA and DNA methylation expression profile, indicating that multiple distinct mechanisms may contribute to AIDS resistance in AGM. Knowledge of the non-pathogenic nature of SIV infection in AGM may provide insight into development of new therapeutic strategies.

## Introduction

Simian immunodeficiency viruses (SIV) belong to the group of lentiviruses that infect non-human primates (NHP). Like HIV-1 and HIV-2, all known SIV subtypes use CD4 as a receptor and either CCR5, CXCR4, CCR2 as a co-receptor [1,2,3]. SIV infection of natural hosts, such as AGM, is usually lack of progression to AIDS despite high viraemia, while HIV infection in humans and experimental SIV infection in rhesus macaques (Macaca mulatta) progress to AIDS. One obvious difference between progressive HIV infection and non-progressive SIV infection is the absence of immune activation during the chronic phase of infection in natural hosts [4].

Low frequencies of CD4^+^ T cells and high frequencies of CD8^dim^ T cells have been shown to exist in healthy adult AGM [5,6]. CD8^dim^ T cells could induce antibody production from B cells in vitro suggesting that CD8^dim^ T cells might supplement for the lack of CD4^+^ T cells in AGM [7]. Down-regulation of CD4 by memory T cells in adult AGM protects these T cells from infection by SIVagm in vivo [8], but the molecular mechanism remains unclear. Epigenetic phenomena are defined as heritable mechanisms that establish and maintain mitotically stable patterns of gene expression without modifying the base sequence of DNA, which include DNA methylation, post-translational histone modifications and RNA-based mechanisms including those controlled by small non-coding RNAs (miRNAs) in mammalian cells [9].

This study aims to identify the differential expressed miRNAs, as well as DNA methylation features, between junior and adult AGM, which may provide insights into AIDS resistance in AGM and add knowledge to development of new therapeutic strategies.

## Materials and Methods

### Ethics Statement

SIVagm-uninfected junior (less than 2 years old) and adult (more than 5 years old) AGM were chosen for analysis. Animals were housed in troop enclosures with an outdoor facility amd fed nonhuman primate chow per day and a combination of fresh bananas, apples, carrots 3 days per week. Toys were available for animals raised for the purpose of environment enrichment. The experiments were conducted in accordance with the recommendations of the ethics provision for experiments on non-human primate of the ethics committee of the Academy of Military Medical Sciences. The protocol was approved by the ethics committee of the Academy of Military Medical Sciences with the identification number 2018032. All protocols were in strict accordance with the recommendations in the Guide for the Care and Use of Laboratory Animals.

### PBMC Isolation

Peripheral blood mononuclear cells were isolated from fresh blood by Ficoll gradient separation. 1×10^7^ cells were mixed with 600 μl mirVana RNA lysis buffer (Ambion, Austin, Texas, USA) to achieve lysis and inactivate endogenous RNAses. Lysates were frozen at −80 °C until further processing.

### RNA isolation and Small RNA Deep Sequencing

Total RNA was isolated using the RNeasy mini kit (Qiagen) according to the manufacturers’ instructions. Small RNA libraries were created using the TruSeq Small RNA Sample Preparation Kit (Illumina) and sequencing performed using HiSeq 2000 sequencer (Illumina). The small RNA deep sequencing and data analysis were carried out by Shanghai KangChen biotechnology company (Shanghai, China).

### Inhibition of endogenous miRNAs

Locked nucleic acid (LNA)-modified anti-miRs (Exiqon) were used for the inhibition of endogenous miRNAs in PBMCs of junior and adult AGM [10]. PBMC were maintained in RPMI1640 supplemented with 100 U/ml penicillin, 100 μg/ml streptomycin and 10% fetal bovine serum (FBS) (Invitrogen). LNA-modified anti-miRs were transfected at a final concentration of 10 nM by Entranster™-R4000 (Engreen Biosystem).

### Real-Time Quantitative Reverse-Transcription Polymerase Chain Reaction

MMLV reverse transcriptase (Takara) was used for the reverse-transcription (RT) reaction and quantitative PCR was performed by an ABI PRISM7500 system (Applied Biosystems). The RT primers and the primer sets specific for each miRNA amplification are shown in Supplementary Table 1. Expression of selected miRNAs and mRNA of the target genes predicted to be targeted by miRNAs inhibited by anti-miR transfection were measured by qPCR using SYBR green chemistry [11]. Primer sequences for the target genes are listed in Supplementary Table 2.

### DNA isolation and MeDIP sequencing

DNA was isolated using the QIAamp DNA mini kit (Qiagen) according to the manufacturers’ instructions. Isolated DNA were fragmented to a size range of ~200-500bp with a Diagenode Bioruptor. About 1 μg of fragmented DNA was prepared for Illumina HiSeq 4000 sequencing as the following steps: 1) End repair of DNA samples; 2) A single ‘A’ base was added to the 3’ ends; 3) Illumina’s genomic adapters were ligated to DNA fragments; 4) DNA fragments were immunoprecipitated by anti-5-methylcytosine antibody; 5) Immunoprecipitated DNA fragments were amplified by PCR amplification; 6) Size selection of ~300-600bp DNA fragments using AMPure XP beads. The completed libraries were quantified by Agilent 2100 Bioanalyzer. The libraries were denatured with 0.1 M NaOH to generate single-stranded DNA molecules, captured on Illumina flow cell, amplified in situ. The libraries were then sequenced on the Illumina HiSeq 4000 following the HiSeq 3000/4000 SBS Kit (300 cycles) protocol.After sequencing images generated, the stages of image analysis and base calling were performed using Off-Line Basecaller software (OLB V1.8). After passing Solexa CHASTITY quality filter, the clean reads were aligned to Chlorocebus-sabaeus genome (ensemble ChlSab1) using HISAT2 software (V2.1.0). Aligned reads were used for peak calling of the MeDIP regions using MACS V2, statistically significant MeDIP enriched regions (peaks) were identified for each sample at a q-value threshold of 10-5 using MACS v2.Statistically significant differentially methylated regions (DMRs) between two groups promoters were identified by diffReps (Cut-off: FC=1.0, p-value=0.05).

### Quantitative SYBR green methylation-specific PCR (QSG-MSP)

QSG-MSP was performed to quantify the levels of CpG DNA methylation of *CD4* and RUNX3.Primers for QSG-MSP were designed using Methyl Primer Express Software v1.0 (Applied Biosystems). Primer sequences for *CD4, RUNX3* and *ACTB* were listed in Supplementary table 3. Quantitative PCR was performed in a 25 μl reaction volume with 12.5 μl of 2×SYBR Green PCR Master Mix (Applied Biosystems), 2.5 pmol of each primer, and 50 ng of bisulfite-treated DNA sample. Thermal cycling was as follows: 95°**C** for 10 min, 40 cycles of 95°**C** for 15 s, 60°**C** for 1 min. The amount of methylated DNA (percentage of methylated reference) was calculated as follows: ratio of quantity of target gene to quantity of target gene of test sample divided quantity of *ACTB*.

### Statistical Analysis

Values were expressed as mean ± SD. The comparative CT method was used in real-time qRT-PCR assay according to the delta-delta CT method. Statistical analyses were performed using GraphPad Prism version 5.01. t test was used to compared statistical differences. The data were considered statistically significant at *P*<0.05.

## Results

### Differential miRNA expression in PBMCs from junior and adult AGM

Different from junior AGM, low frequencies of CD4^+^ T cells and high frequencies of CD8^dim^ T cells were observed in healthy adult AGM, which may partially explain the absence of SIV-induced disease progression. To investigate the role of miRNA in this process, we profiled the PBMC miRNA expression in 3 junior AGM and 3 adult AGM. As shown in Figure 1a, 3 miRNAs were up-regulated while 15 miRNAs were down-regulated with over 1.5-fold change in adult AGM (*P*<0.05). To validate the profiling data, miRNAs in Table 1 were further analyzed via quantitative RT-PCR between PBMCs from 5 junior AGM and 5 adult AGM. Results showed a 3.6-fold decrease for hsa-miR-215-5p (*P*<0.01), a 2.0~2.5-fold decrease for hsa-miR-194-5p, hsa-miR-99b-5p, hsa-miR-125a-5p (all *P*<0.01), a 2.0~2.5-fold increase for hsa-miR-95-3p (*P*<0.01), a 1.5~2.0-fold decrease for hsa-miR-126-3p, hsa-miR-181b-5p, hsa-miR-140-5p, hsa-miR-199a-3p, hsa-miR-10a-5p (all *P*<0.01), and a 1.5~2.0-fold increase for hsa-miR-151a-3p, hsa-miR-10b-5p (all *P*<0.05) (Figure 1b and 1c).

**Figure 1.**
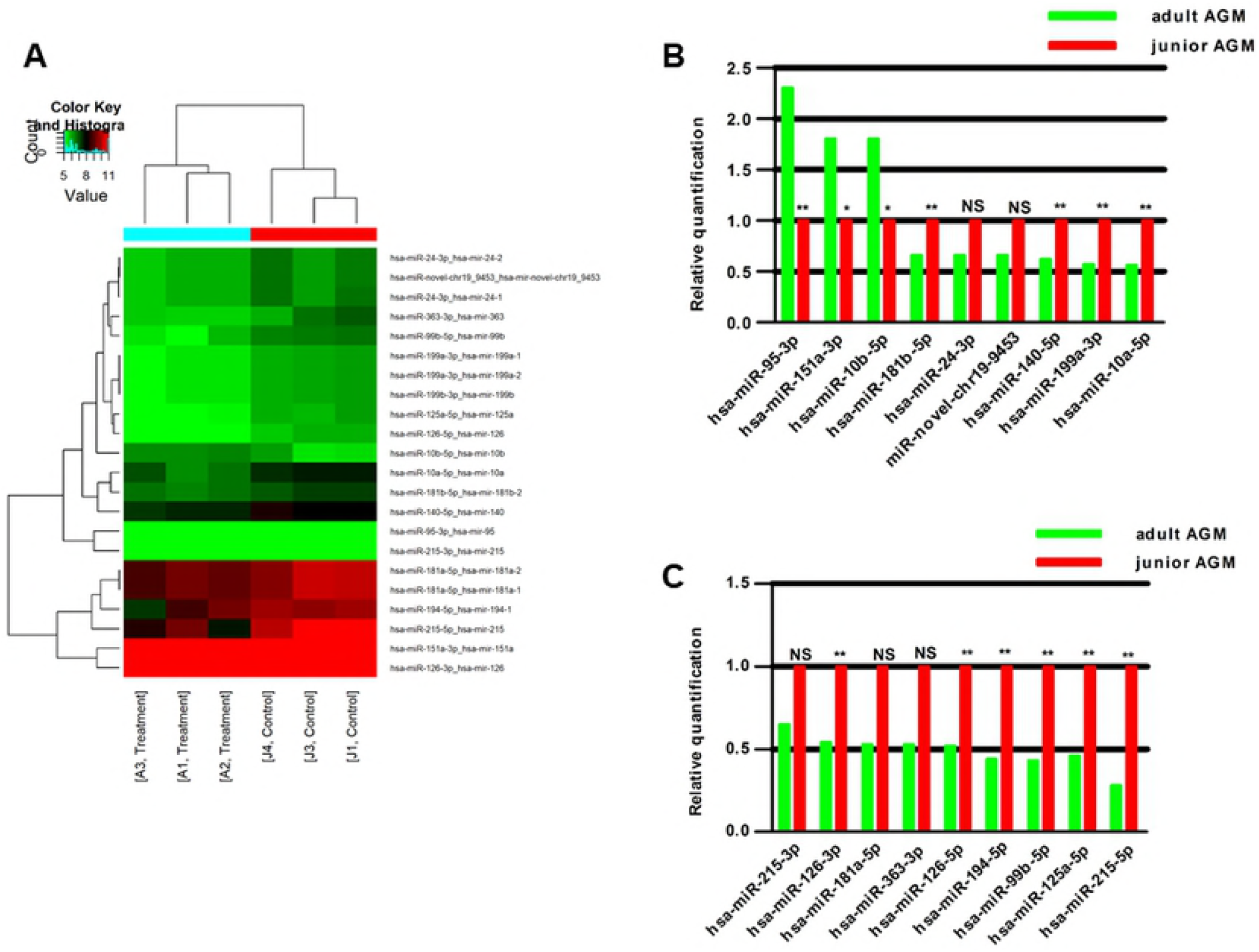
miRNA profiling and validation of junior and adult AGM PBMCs. (**A**) Heat map of miRNA microarray expression data from PBMC samples of junior AGM (n=3) and adult AGM (n=3). (**B&C**) Validation of miRNA microarray data by quantitative reverse-transcription polymerase chain reaction. The relative expression of miRNAs was normalized to expression of the internal control (U6). The *P* values were calculated by 2-sided Student *t* test. \**P*<0.05; \*\**P*<0.01.

### Effects of selected miRNAs on AGM *CD4*, *CD8α* and *MAZR*

As MAZR is a protein suppressor of the *CD8α* enhancer region [12], we investigate whether the above miRNAs target AGM *CD4, CD8α* and *MAZR*. hsa-miR-151a-3p was predicted to target AGM *CD4* and *MAZR* while hsa-miR-140-5p, hsa-miR-126-3p and hsa-miR-194-5p were predicted to target AGM *CD8α* (Figure 2a).Then, the potential miRNA-target pairs were examined by inhibiting these endogenous miRNAs in PBMCs using LNA-modified anti-miRs. Anti-miR treatment may cause the increase on the abundance of the target mRNA if it is indeed suppressed by the endogenous miRNA via mRNA degradation. As shown in Figure 2b, the selected miRNAs were all validated to target AGM *CD4, CD8α* and *MAZR*, respectively. Interestingly, none of these miRNAs possess target sites in cynomolgus macaque (CM) *CD4, CD8α* and *MAZR*, which may partially explain the differences in AIDS resistance between these two species.

**Figure 2.**
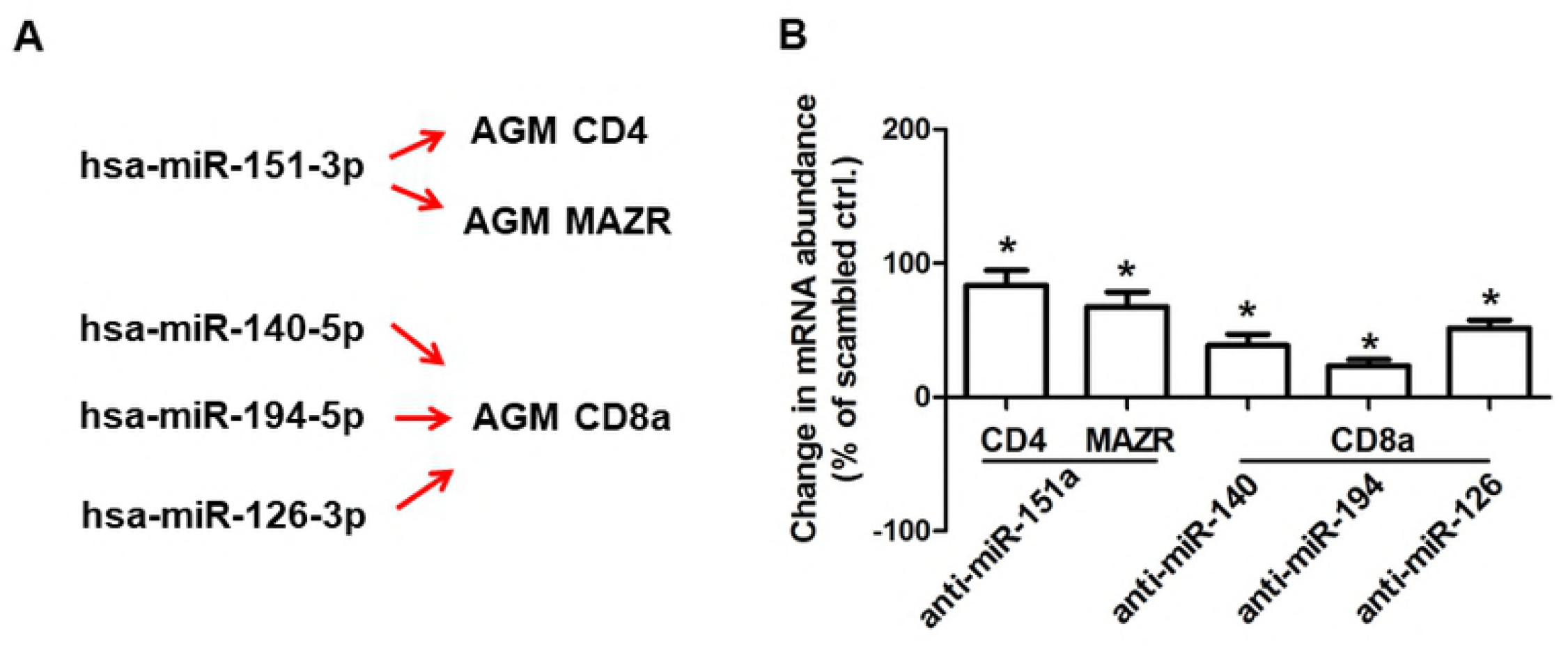
Summary of miRNA-target pairs supported by the anti-miR transfection experiment. The mRNA abundance for the predicted targets was significantly increased in PBMC after inhibition of the indicated endogenous miRNA. The *P* values were calculated by 2-sided Student *t* test. \**P*<0.05.

### Differential DNA methylation in PBMCs from junior and adult AGM

Besides miRNA profiles, we wondered whether differences also exist in DNA methylation profiles and performed MeDIP sequencing. Genes with significantly different DNA methylation signals in promoter regions were selected and the Top10 biological processes (BP) were listes through Gene Ontology (GO) analysis (Figure 3). RUNX3 is required for the establishment of epigenetic silencing of CD4 in cytotoxic-lineage thymocytes [13]. Our results showed hypomethylation in *RUNX3* promoter and up-regulation of its mRNA in adult AGM (Figure 4b and 4f), as well as hypermethylation in CD4 promoter (Figure 4a), which together may account for the CD4 repression in adult AGM T cells. Although CD8α was over-expressed in adult AGM T cells, we did not find similar DNA methylation changes in proteins regulating CD8 expression, such as MAZR, TOX, Ikaros [14], suggesting other mechanisms may exist. As a coreceptor used by SIV *in vivo*, the level of CCR5 is very low on CD4^+^ T cells of natural host species [15,16]. CXCR6 is another potential alternative coreceptor for SIV as an efficient entry pathway in vitro [17,18]. Analysis through QSG-MSP demonstrated both hypermethylation in CXCR6 and CCR5 promoters (Figure 4c and 4d) and both mRNAs were down-regulated in adult AGM PBMCs (Figure 4f), providing another insight into AIDS resistance in African green monkey. Recently, the genome of another natural host, sooty mangabeys (*Cercocebus atys*), has been sequenced and assembled [19]. Also, the *C.atys* immune-regulatory protein intercellular adhesion molecule 2 (ICAM-2) was found to possess a major structural change in exons 3-4 and expression of this variant leads to reduced cell surface expression of ICAM-2. Our results showed that hypomethylation in promoter region of ICAM-2 existed in adult AGM (Figure 4e) and expression of ICAM-2 was significantly up-regulated in adult AGM (Figure 4f). Furthermore, hypomethylation in the promoter region of *PTK2*, as well as hypermethylation in the promoter region of *WWP2* was demonstrated in adult AGM (Figure 5a and 5b), in which hsa-miR-151a-3p and hsa-miR-140-5p was located, respectively. This indicates a cross-link between roles of DNA methylation and miRNA expression in AIDS resistance in AGM.

**Figure 3.**
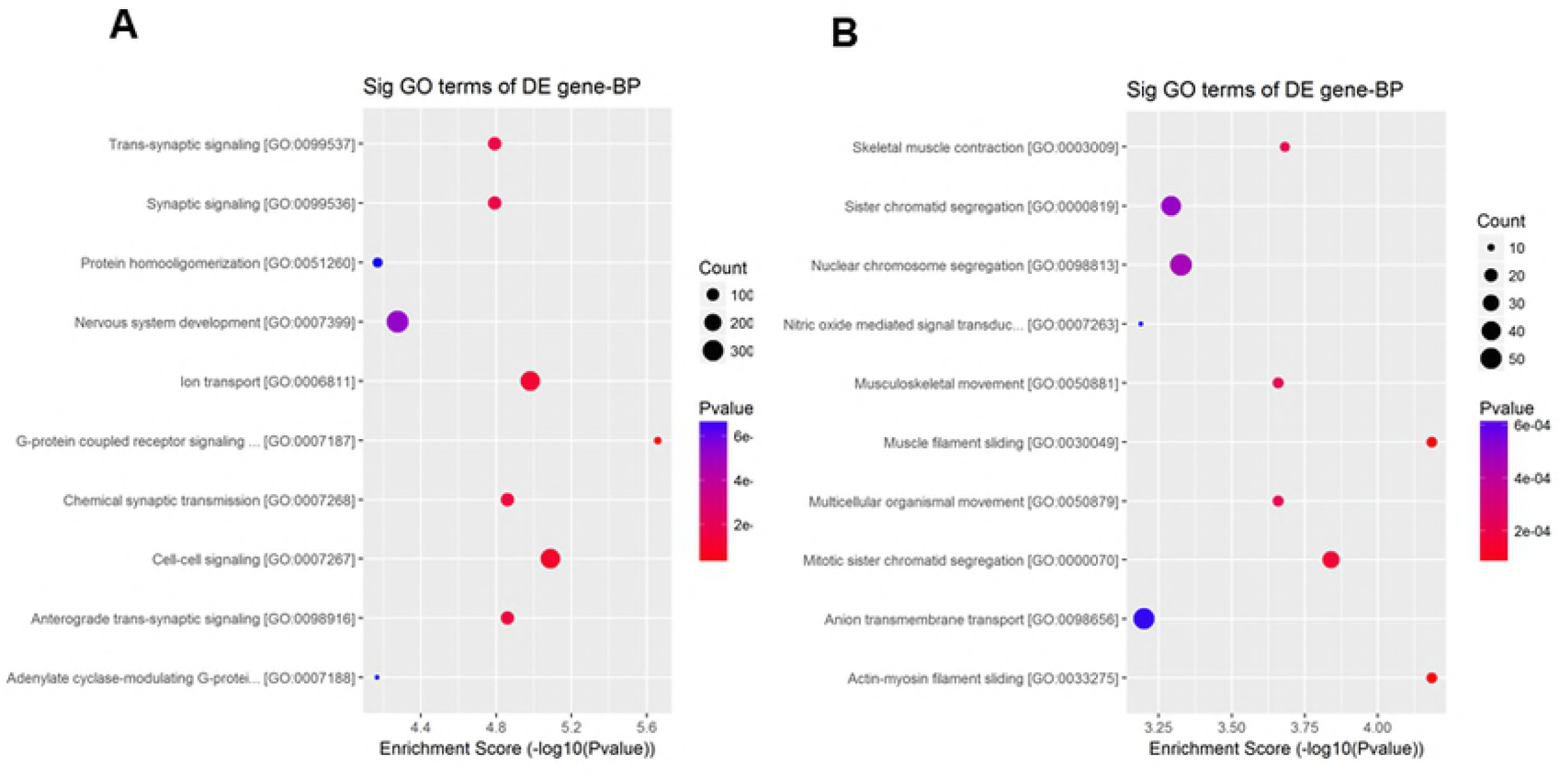
Enrichment map of GO categories for biological processes. A indicates those hypermethylated genes in adult AGM while B indicates those hypomethylated genes. Colors represent *P* values on a log scale (with red corresponding to the most highly significant, *P*<0.05). Node size represents the number of genes in a category.

**Figure 4.**
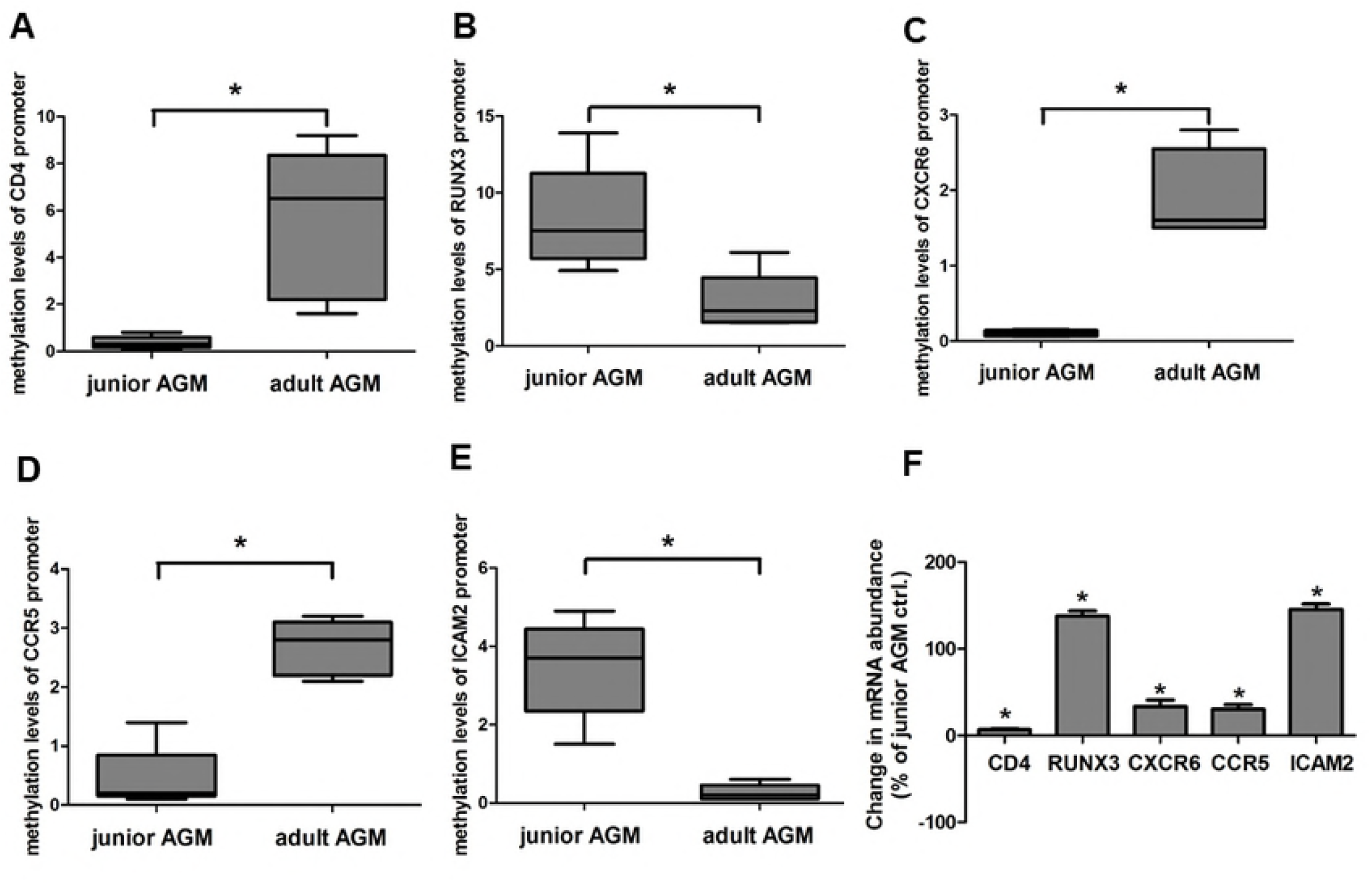
Comparative analysis of differential DNA methylation between junior and adult AGM. (**A,B,C,D&E**) Quantitative analysis of the levels of CpG DNA methylation of *CD4*, *RUNX3*, *CXCR6*, *CCR5* and *ICAM2*. (**f**) Quantitative RT-PCR analysis for mRNA expression of the target genes. The *P* values were calculated by 2-sided Student *t* test. * stands for *P*<0.05.

**Figure 5.**
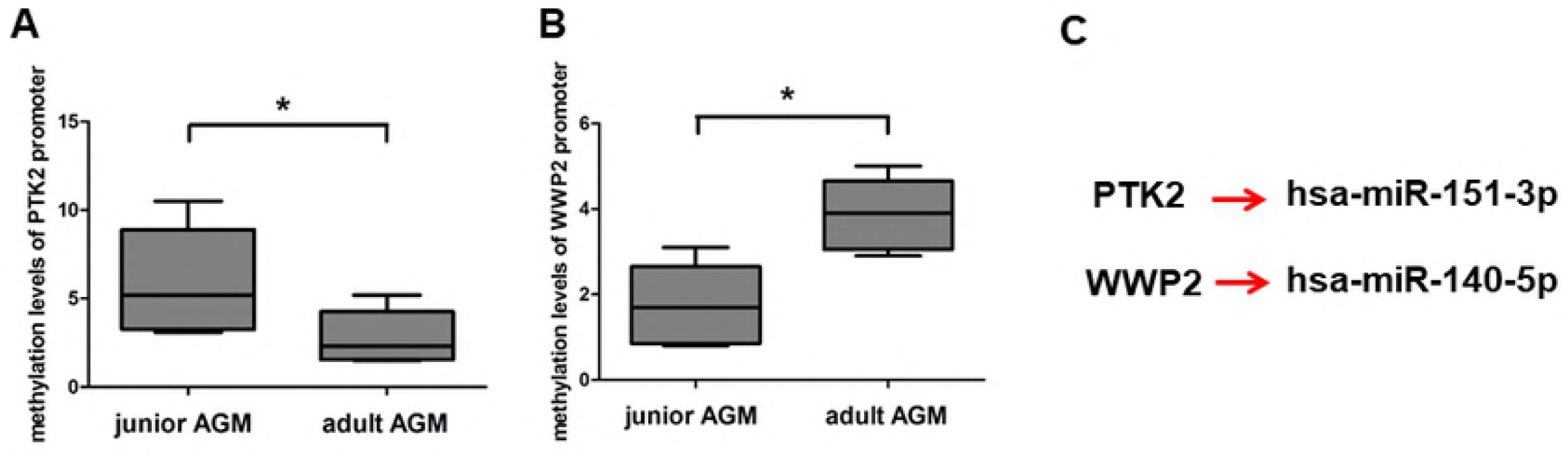
Quantitative analysis of the levels of CpG DNA methylation of PTK2 and WWP2. (**A&B**) Quantitative analysis of the levels of CpG DNA methylation of PTK2 and WWP2. (**C**) Schematic show of the cross-link between DNA methylation and miRNA expression.

## Discussion

Natural hosts have co-evolved with SIV and are capable of avoiding disease progression, of which mechanisms may diverge. Unlike AGM, sooty mangabey (SM) maintains healthy frequencied of CD4^+^ T cells and genome sequencing has identified two gene products (ICAM-2 and TLR-4), which show structural differences that may influence cell-surface expression (ICAM-2) and downstream signalling (TLR-4) [19]. In AGM, the co-evolution with SIVagm may be accounted for partially by the development of CD4^−^CD8a^dim^ T cells from memory CD4^+^ T cells. Like other natural hosts of SIV, CD4-like immunological functions can be elicited by CD4^−^ T cells in AGM [20] and preservation of CD4^+^ T cell function may contribute to the lack of immune activation in AGM and SM [5,21,22].

It has been shown that memory CD4^+^ T cells down-regulate CD4 and up-regulate CD8α [8], but the actual mechanism(s) underlying the switch from CD4^+^ to CD4^−^CD8a^dim^ remains unclear. Previous studies have demonstrated genetic differences between certain regulatory elements from AGM compared to other primates [23]. miRNAs result in translational suppression of suppression of target mRNA in all known animal and plant genomes [24]. DNA methylation is an epigenetic modification typically associated with stable transcriptional silencing and plays an important role in several biological processes associated with development and disease [25,26]. In this study, the miRNA expression patterns were found to be associated with the switch from CD4^+^ to CD4^−^CD8a^dim^ in adult AGM. The up-regulated hsa-miR-151a-3p was shown to target both AGM *CD4* and *MAZR*, while the down-regulated hsa-miR-140-5p, hsa-miR-126-3p and hsa-miR-194-5p were shown to target AGM *CD8α*. And none of these miRNAs possess target sites in cynomolgus macaque (CM) *CD4, CD8α* and *MAZR* reflecting differences in AIDS resistance between these two species. Differential DNA methylation in promoter regions of *CD4, RUNX3, CXCR6, CCR5, ICAM-2*, as well as *PTK2* and *WWP2*, was also demonstrated indicating that multiple distinct mechanisms may contribute to AIDS resistance in AGM. Knowledge of the non-pathogenic nature of SIV infection in AGM may provide insight into development of new therapeutic strategies.

## ABBREVIATIONS

AIDS,: acquired immune deficiency syndrome;
MAZR,: Myc-associated Zn finger-related factor;
MeDIP,: methylation DNA immunoprecipitation;
CXCR6,: C-X-C motif chemokine receptor 6;
CCR5,: C-C motif chemokine receptor 5;
RUNX3,: Runt-related transcription factor 3;
ICAM2,: intercellular cell adhesion molecule-2;
CCR2,: C-C motif chemokine receptor 2;
MMLV,: moloney murine leukemia virus;
ACTB,: actin β;
TLR-4,: Toll-like receptor 4;
PTK2,: protein tyrosine kinase 2;
WWP2,: ww domain-containing protein 2.

## Acknowledgments

The authors are grateful to Professor Jun-feng Li (Institute of Microbiology and Epidemiology, Beijing, China) for technical support in QSG-MSP.

## Author Contributions

Conceived and designed the experiments: XZ ZY. Performed the experiments: XZ JW. Wrote the paper: XZ.

## Supporting Information Legends

**Table 1. Properties of miRNAs differentially expressed in PBMCs of adult and junior African green monkeys**.

**Supplementary Table 1. Sequences of the primers used in the SYBR-green-based quantitative RT-PCR validation**.

**Supplementary table 2. Primer Sequences for quantitative analysis of AGM CD4,CD8a, MAZR**.

**Supplementary Table 3. Primer Sequences and PCR Conditions for MSP Analysis**.

